# Action in auctions: neural and computational mechanisms of bidding behavior

**DOI:** 10.1101/464925

**Authors:** Mario Martinez-Saito, Rodion Konovalov, Michael A. Piradov, Anna Shestakova, Boris Gutkin, Vasily Klucharev

## Abstract

Competition for resources is a fundamental characteristic of evolution. Auctions have been widely used to model competition of individuals for resources, and bidding behavior plays a major role in social competition. Yet how humans learn to bid efficiently remains an open question. We used model-based neuroimaging to investigate the neural mechanisms of bidding behavior under different types of competition. Twenty-seven subjects (nine male) played a prototypical bidding game: a double action, with three “market” types, which differed in the number of competitors. We compared different computational learning models of bidding: directional learning models (DL), where the model bid is “nudged” depending on whether it was accepted or rejected, along with standard reinforcement learning models (RL). We found that DL fit the behavior best and resulted in higher payoffs. We found the binary learning signal associated with DL to be represented by neural activity in the striatum distinctly posterior to a weaker reward prediction error signal. We posited that DL is an efficient heuristic for valuation when the action (bid) space is continuous. Indeed, we found that the posterior parietal cortex represents the continuous action-space of the task, and the frontopolar prefrontal cortex distinguishes among conditions of social competition. Based on our findings we proposed a conceptual model that accounts for a sequence of processes that are required to perform successful and flexible bidding under different types of competition.

## Introduction

We often deal with situations where buyers and sellers meet to exchange goods at prices determined by fluctuations in supply and demand. Perceived market competition influences human bidding (Fischbacher et al., 2009; van den Bos et al., 2008) and even the value of commodities traded by non-human animals. For instance, baboons (Henzi & Barrett, 2002) and vervet monkeys (Fruteau et al., 2009) demonstrate the effect of market competition on the price of natural currencies such as food or grooming. Indeed, biological auctions are used to model competition between species and individuals (Reiter et al., 2015). Despite its key importance in social behavior and financial modeling, the neural mechanisms of decision-making under market competition are still unclear. In particular, how do we learn bidding strategies across different market scenarios? Here, we investigate the neural mechanisms underlying bidding under different conditions of competition.

The study of bidding behavior lies at the intersection of behavioral economics, game theory, and cognitive neuroscience. Much previous research has focused on simple sequential game theoretic paradigms, such as the ultimatum game (UG; Güth et al., 1982; Sanfey et al., 2003). Behavioral studies have shown that competition in UGs among proposers leads to higher bid offers (Roth et al., 1991), and in general it pushes players towards Nash equilibria with tell-tale lower rejection rates (Fischbacher et al., 2009). A combination of fairness concerns and decision errors has been put forward to explain the effect of competition on offer distributions in UGs (Fischbacher et al., 2009), but it is not clear how offers are picked in more general settings. In simultaneous bidding paradigms, Quantal Response Equilibrium (McKelvey & Palfrey, 1995), a normative solution concept from game theory, has been shown to capture behavior well. However, this model offers little insight about biological learning mechanisms and requires costly computations based on beliefs about other players. In repeated games, players typically demonstrate an extended adaptation to the environment’s conditions (Fudenberg & Levine, 1998; Grosskopf, 2003; Roth et al., 1991), and very simple models have been shown to perform robustly as long as enough information about other players is provided (Fudenberg & Levine, 2009). Moreover, behavioral economics experiments show that adaptive learning algorithms explain bargaining behavior well (Camerer & Ho, 1999; Erev & Roth, 1998; Mookherjee & Sopher, 1994). Thus, a parsimonious learning model should be suitable for explaining offer distributions under changing supply and demand conditions.

Previous neuroimaging studies investigated bargaining games, but focused on strategic deception and uncertainty about trustworthiness (Bhatt et al., 2010, 2012) or examined the influence of loss contemplation under social contexts in overbidding (Delgado et al., 2008). In this study, we investigated the neural mechanism of bidding behavior under different conditions of competition. Subjects played the role of buyers in a double auction in three different market types, which differed in levels of supply and demand. To investigate buyer’s decisions, we set the transaction price to equal the buyer’s bid, which in case of acceptance becomes the final price, while rejection was set to be the worst outcome. This paradigm is similar to online auctions such as eBay auction, where multiple buyers bid for a good, and in financial transactions with buy limit orders (assuming that buyers are strongly incentivized to acquire the good). In these scenarios, repeated bidding serves to ‘probe’ the market and estimate its current clearing price in a trial-and-error fashion, and thereby the buyer learns to bid more efficiently given the estimated clearing price and her needs.

Although traditionally theoretical accounts of adaptive learning in decision-making tend to focus on model-free reinforcement learning (RL), algorithms that are beyond this minimal account may be more suitable for bidding. One such framework that is particularly suitable for bidding, directional learning (DL), suggests a simple adaptive strategy that takes into account that the available bids are ordered consistently (Selten & Buchta, 1994) and requires a representation of a one-dimensional continuum. According to DL, profitable bids exhibit a simple Markovian dependence on the immediately previous outcome: it is adjusted up (down) if it was too low (high) in the previous period.

To our knowledge, DL models have not been used in neuroimaging studies to probe the neural correlates of economic decision-making. However, numerous functional magnetic resonance imaging (fMRI) studies have shown that RL operational variables, such as expected value and reward prediction error (RPE), can be used to trace neural correlates of adaptive learning (e.g. Montague et al., 2006; Ruff & Fehr, 2014). For example, neural correlates of RPE have repeatedly been located in the ventromedial prefrontal cortex (vmPFC) and the ventral striatum (Bartra et al., 2013; O’Doherty et al., 2004). But such studies often use relatively simple decision-making tasks, structured specifically to be solvable by RL in a reasonable time, often with discrete response policies, while economic tasks involving continuous decision variables and policies that need to be structured over such real-value scales have been explored to a lesser extent. Here, we focus specifically on the neural underpinnings of DL and RL strategies that drive repeated bidding behavior under different types of buyer/seller competition.

## Materials and Methods

### Subjects

Twenty-seven subjects (nine males, two left-handed, after discarding two of the initial 29 subjects due to excessive head motion) took part in the experiment. All subjects were queried to exclude histories of neurological pathologies. After a briefing, all subjects gave informed written consent. The protocol was approved by the local university’s ethics committee.

### The double auction paradigm

To probe neural mechanisms of bid-learning, we used a modified version of the double auction is a standard paradigm in multiplayer game theory where players try to maximize their respective benefit by means of a single-shot transaction (Fudenberg & Tirole, 1991). Subjects played the role of buyers in a double auction with first-price sealed bids, and with opponents assigned by repeated random matching, in three different market types (Figure 1A).

**Figure 1.**
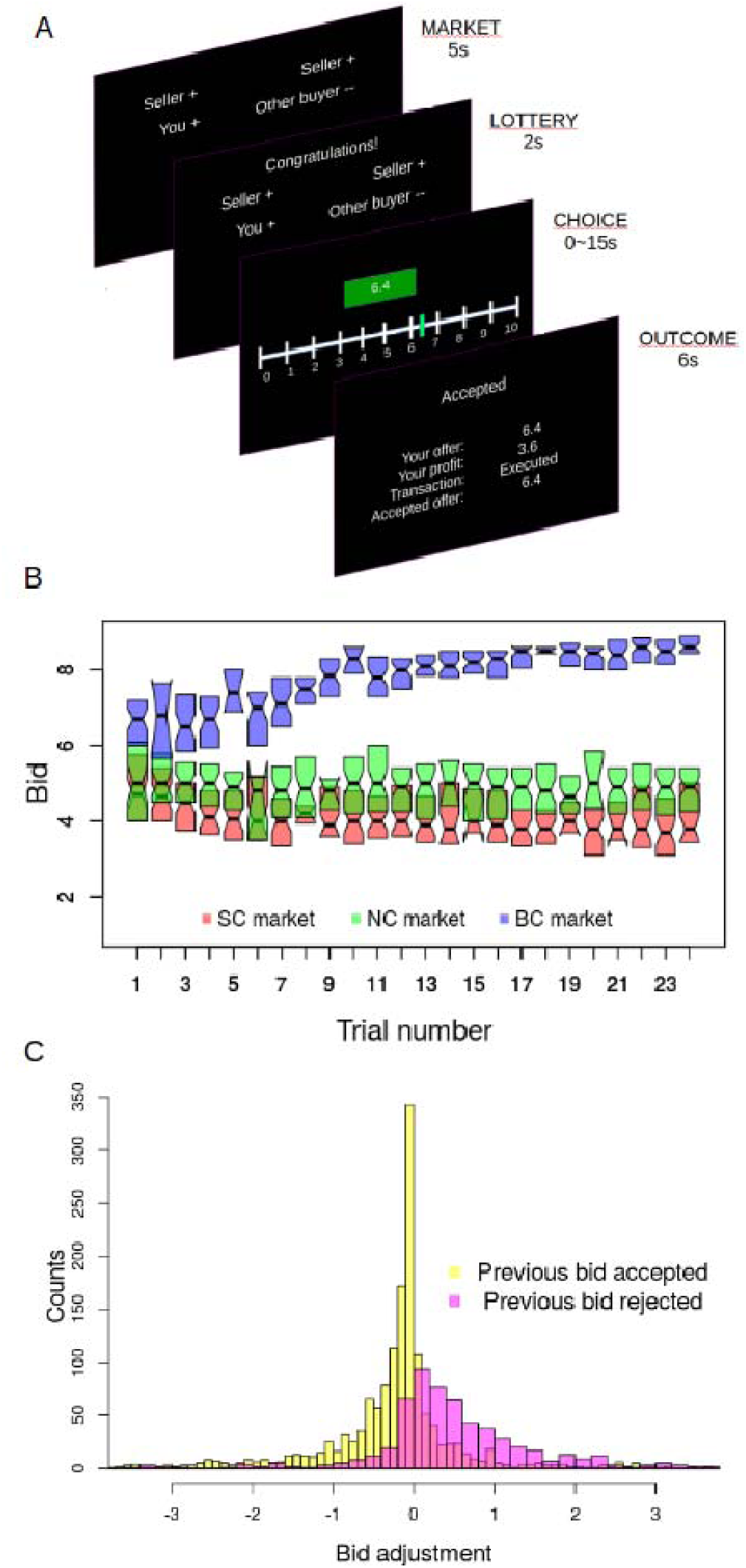
Task design and behavioral results. (A) Each trial consisted of four stages: market type announcement, lottery, bid selection, and game outcome feedback. During the market announcement stage (MARKET), the subject was informed of the market type of the current trial. The next stage (LOTTERY) indicated whether the subject would go forth to the next stage or be redirected to the beginning of the next trial. In the former case, a Likert scale was displayed, and the subject had to choose her bid by sliding a vertical bar (CHOICE). Finally, the game outcome stage (OUTCOME) signaled whether the bid was accepted (ACCEPTED) or rejected (REJECTED). (B) Behavioral learning dynamics of bids across all subjects. Box “hinges” represent the first and third quartiles. (C) Bid adjustments were contingent on the previous trial’s outcome of the same market type.

The three market types differed in the number of sellers and buyers. In the *seller competition market* (SC), there were two sellers and one buyer (the subject); in the *no competition market* (NC), one seller and one buyer (the subject); and in the *buyer competition market* (BC), one seller and two buyers (one of them being the subject). In all market types, the outcome of the transaction was determined by pitting the highest buyer’s bid against the lowest seller’s ask price. If the former was strictly lower than the latter, then the transaction was not consummated, and the subject received the disagreement outcome: zero monetary units (MU). Otherwise, the subject received *10*-*b* MU, where *b* is the bid of the subject. Hence the win/loose structure was asymmetric: the win from an accepted bid was dependent on the bid amount, while the loss of fixed at 10 MU. We focused exclusively on buyer behavior, unlike previous studies analyzing all players’ behavior (Grosskopf, 2003; Güth et al., 1982). The clearing price was set to be the maximum bid in order to study buyer behavior specifically.

### Task description

Subjects were informed that they were participating in a game investigating decision-making. The game paradigm required buyers to fix their bids in advance. Their task was to buy fish on a market using a 10-point Likert scale with increments of 0.1 MU. The initial position of the cursor on the Likert scale was randomized across trials. Collected fish led to a payoff: *p* = *10* – *b*, where *b* was the bid value in task MU and 10 represented the maximum endowment the player could make use of in every transaction. Opponents were prerecorded human subjects replayed by a computer. In each trial, subjects played in one of the market types, which were looped throughout the experiment (24 blocks of 3 market types) in the order determined by a fixed sequence without repetition (of SC, NC, and BC). One of the six possible sequences was pseudo-randomly and independently assigned to each subject.

At the beginning of a trial, a MARKET stage (duration=5s, Figure 1A) informed subjects of the market type in the current trial. Next, a LOTTERY (duration=2s) stage consisted of a lottery determining whether subjects would be allowed to enter the market or not. In one of every six trials, subjects were not allowed to enter the market and had to move to the next trial. Otherwise, subjects entered the market and the CHOICE stage started. During the CHOICE stage (self-paced, but with a prompt to answer quicker after 15s), subjects had to purchase (by bidding) fish in a market using a 101-point slider scale. The feedback screen (OUTCOME, duration=6s) displayed the outcome of the transaction and the profit earned. In BC trials, when the competitor outbid the subject, that bid was made visible to the subject. Sellers’ ask prices were never disclosed. All inter-stimulus intervals were jittered between 5 and 7s following a uniform distribution of duration 2s. The LOTTERY stage was included to assess the subject’s differential neural response to being rejected from each market type. However, we found no significant differences in this respect.

Every subject played 24 trials of each market type (72 in total). The duration of each trial depended on the bid selection time and ranged from 21s to 61s, with an average of 39s. The total duration of the experiment was approximately 50 min.

The instructions explicitly informed subjects that they would play against prerecorded human players who had played the same game before against other human opponents. Our design precluded subjects from trying to manipulate their opponents’ behavior in a sophisticated manner (Camerer et al., 2002; Bhatt et al., 2010). In each trial, the actions of the subjects’ opponents were matched according to the trial order of each market type (repeated random matching). Once inside the scanner but before the scanning started, subjects were trained on 6–10 trials, encompassing all market types (at least two trials of each market type). The training phase ended after subjects successfully and consistently manipulated the button box by placing their intended bid and then reported understanding the task.

After scanning, subjects were rewarded according to the following reward scheme (Roth et al., 1991): a fixed compensation of 300 Russian rubles (~5USD) for participation, in addition to a bonus equal to the sum of the profit earned in three random trials multiplied by 15 MU (~5–12 USD in total).

The prerecorded data were recycled from a previous pilot study that implemented the same paradigm. Its design was identical to that of the present study with the following exceptions: 32 subjects played with real opponents in anonymous groups on desktop computers with conventional keyboards, and they played against each other, simultaneously, in the same room. The game was programmed in z-Tree (Fischbacher, 2007). Subject roles were randomly assigned to buyer *or* seller throughout the duration of the experiment. Both seller and buyer had to set their respective ask prices and bids beforehand. The total number of trials amounted to 240 (40 periods with 6 rounds per period).

Our task is a one-shot game because opponents are assigned by repeated random matching. However, given that subjects play repeatedly in the same three market types, this task also displays attributes of sequential games in the sense that what is being learned is not the type of one opponent, but the behavior of a population of players as a whole. This topic has been previously explored from the viewpoint of strategic teaching (Camerer et al., 2002).

### Stimulus presentation and response collection

The visual stimuli were projected with an LCD projector onto a rear screen. This screen was reflected by a mirror attached to the MRI head coil, subtending approximately 20 degrees of visual angle. The task was programmed using Presentation software (version 18.0, Neurobehavioral Systems, USA). Responses were collected through three response buttons: the right thumb shifted the cursor to the right, the right index shifted it to the left, and the left thumb confirmed bid choices.

### Computational algorithms of adaptive learning

We implemented, fitted, tested, and simulated six learning algorithms, including model-free RL and model-based DL algorithms with ad-hoc parameters (Table 1). The dataset consisted of the aggregated sequence of all trials played by the 27 subjects with the same prerecorded opponents. The null algorithm, consisting of assigning uniform probability to all outcomes, was run as benchmark. The important parameters were the learning rate (a measure of how much weight was given to recent feedback with respect to older feedback) and the randomness of choice, embodied in the inverse temperature of the softmax function (a measure of degree of action selection randomness) for RL algorithms, and in the dispersion parameters for DL algorithms. The dispersion parameters could be specific to the upper or lower side of the preferred bid, and to the previous trial outcome contingency.

**Table 1.**
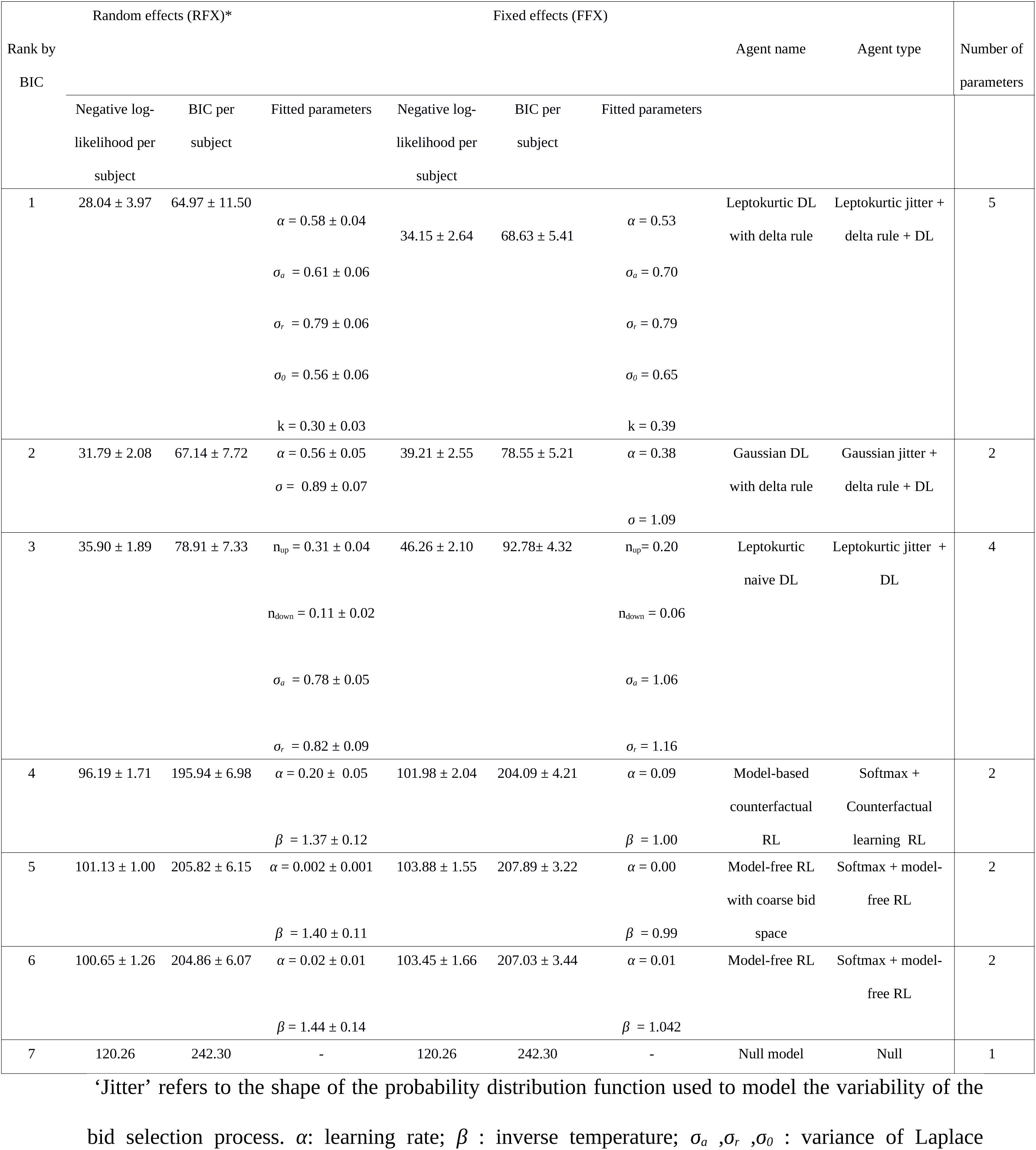
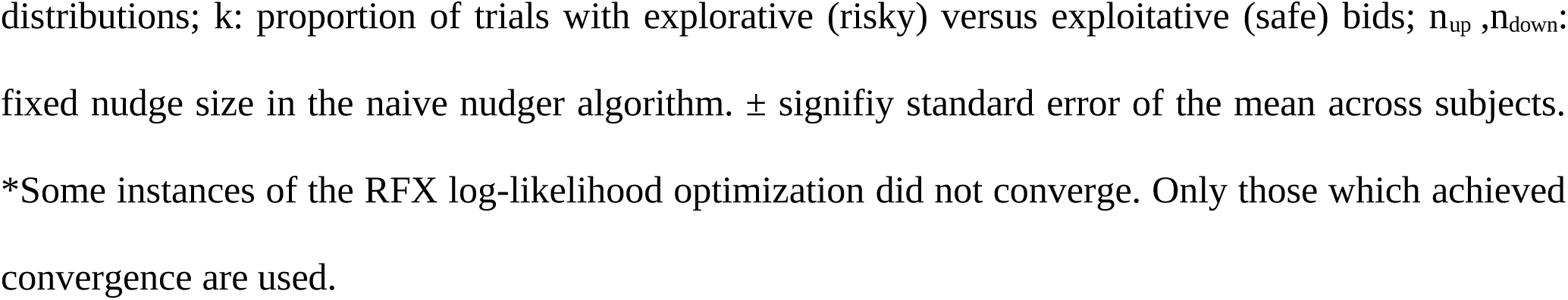
Ranks and BIC scores for all fitted algorithms.

In our task, there is only one state (each of the market types), unlike typical scenarios for RL agents, where the phase space comprises many states. The “native” action space consisted of 101 bid sizes. Although schemes for RL on continuous spaces have been proposed (Doya, 2000; van Hasselt & Wiering, 2007), we opted to use a coarse “binned” representation of the native action space for our RL models, fitting multiple candidate algorithms informed by task-specific assumptions. For the DL algorithms we used the native action space.

To design the computational learning algorithms, based on preliminary data and heuristic reasoning, we devised a conceptual learning model of repeated bidding. The model requires at least three computational processes: (a) recognition of the different market types, (b) an internal representation of bid space, and (c) model-based learning optimizing bid choices.

#### Model-free RL

First, we modeled participants’ decisions using a model-free RL algorithm which learned to ascribe, maintain, and update values attached to actions (Sutton & Barto, 1998). Here the problem lies solely in choosing a single bid repeatedly. The basic action-value updating equation was

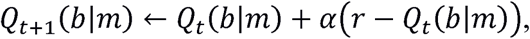

where *Q(b*|*m)* is the action-value function with a value for each possible bid *b* given market type *m* at trial *t*, and *α* is the learning rate regulating the speed of action-value updating. Action values were learned independently for each of the three market types. The policy for selecting a bid in each trial was a conventional softmax function

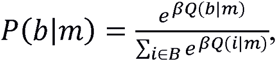

where P(*b*|*m*) is the probability of choosing bid *b* in market type *m*, *β* is the inverse temperature parameter regulating the randomness in action selection, and *B* is the space of actions (bids). Clearly such naive algorithm would perform very poorly given that it neglects the incentive structure of the game and the low ratio of trials to possible actions. Therefore, we binned the 101 actions into 11 uniform tiles (which speeds up learning), and we initialized the action value function distribution for each market type with a Beta distribution (with the two usual parameters and one scaling parameter) fit to the subject-pooled first trial bids (which furnishes efficient priors based on the subject’s pre-game beliefs for choosing bids).

#### Model-based RL with counterfactual learning

Other models are more suitable when relevant prior information is known about the task structure that can be crucial to solve complex tasks where model-free RL becomes unwieldy. We used counterfactual learning, an extension of model-free RL where the value function is updated contingent not only on the currently chosen action feedback, but also on non-chosen actions based on a model about the contingent rewards of foregone actions. This model is derived from the observation that in auctions, any bid lower than the ask price of the seller (and thus lower any previously accepted bid), would have been also accepted, had it been chosen. Value updating occurs for actions that were not chosen, but which are nevertheless updated based on the assumption that they would have been updated had they been chosen. Here, counterfactual learning is carried into effect explicitly as a model-based RL algorithm which asymmetrically updates (through a delta rule) the whole domain of bid choices every time a bid is selected, conditional on both the bid value and the feedback. Overall, it can be considered a hybrid of value-function and model-based algorithm.

We applied the following rule sketch: for every bid *b* selected, if it is accepted (rejected), increase (decrease) the value of the action-value function for all *a*>*b*. This however does not specify how much to decrease or increase the value, and for which actions. We chose to update values conditioned on the outcome of the current transaction only for the higher or lower range of bids for accepted and rejected trials respectively, as follows.

If accept: For all i < b, *Q*_*t*+1_(*i*|*b*, *m*) → *Q_t_*(*i*|*b*, *m*)

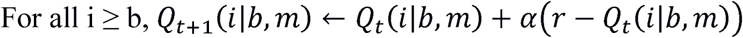

If reject: For all i ≤ b, *Q*_*t*+1_(*i*|*b*, *m*) → *Q_t_*(*i*|*b*, *m*) + α(0 – *Q_t_*(*i*|*b*, *m*))

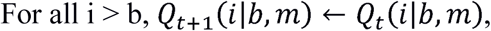

where α is the learning rate and *r_i_* is the counterfactual reward, that is, the reward the player would have received had she selected the bid *i*. For the current trial bid *b*, *r_i_* = *r_b_* = *r*, the reward actually obtained. The action value function distribution was initialized for each market type with a Beta distribution fit to the pooled first trial bids.

#### DL as value-free, model-based learning

DL is a learning mechanism suggested for repeated games (Selten & Buchta, 1994). DL is suitable only under specific circumstances: the space of feasible actions should be a totally ordered set (actions should represent some magnitude varying monotonically), and there should exist a unique optimal action, so DL implies having an a priori knowledge about the structure of the environment. Our task, the double auction, satisfies these conditions, since assuming that the seller holds a given ask price, there is a unique minimal bid below which all bids are rejected. DL is effectively a myopic policy that operates without the need of action-value functions, by nudging the bids up or down depending on a directional signature (DS): whether the previous bid was accepted or rejected. However, the payoff structure of choices around the optimal action is markedly asymmetric in our study, since overbidding entails a reduction of the profit proportional to the overbid, but underbidding entails a null profit.

In every trial of market type *m*, DL is implemented by picking a bid from a unimodal probability distribution *P(b*|*m)* centered in the preferred bid (the lowest accepted bid estimate). If the selected bid is accepted (rejected), then the preferred bid is increased (decreased). The preferred bid for the first trial of each market type was set to equal the mean of the pooled first trial bids. Unlike RL algorithms, DL algorithms lack the notion of expected value, and therefore of RPE. However, it is possible to defined a *pseudo*-*RPE signal* as a RPE where the expected value is assumed to be the currently preferred bid. This framework still leaves unspecified how much to decrease or increase the preferred bid, so we devised and fitted three adaptive learning algorithms based on DL.

#### DL delta rule with Gaussian noise

This is perhaps the simplest conceivable DL model. We can update values conditioned on the outcome of the current trial by making the gain depend on the value of the current preferred bid and the reward received: *A*_*t*+1_(*m*) ← *A_t_*(*m*) + *α*(*r* – *A_t_*(*m*)), where *α* is a gain akin to the learning rate in RL, *A_t_* is the preferred bid at trial *t*, *m* is the market type (SC, NC, or BC), and *r* is the reward. Here, the policy for bid selection accounts for noisy decision-making by means of a Gaussian distribution function of bids around the preferred bid: 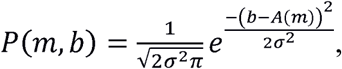 where *σ* is the standard deviation and *A(m)*, which is equal to the preferred bid for market type *m*, is the mean.

#### Naive DL with asymmetric leptokurtic noise

This algorithm consists of simply “nudging” the bid up and down but taking into account the incentive structure of the game by doing it asymmetrically with respect to the two sides of the preferred bid. Contingent on the outcome of the transaction, the preferred bid is updated as follows:

If accepted, *A*_*t*+1_(*m*) → *A_t_*(*m*) – *n_down_*, and if rejected,.

We chose ad hoc a leptokurtic probability distribution function to model the noise around the preferred bid because it fits the data better than the Gaussian distribution (see Figure 1C). The distribution of bids (Figure 1C) is markedly asymmetric and non-Gaussian, specifically with fatter tails and a thinner peak.

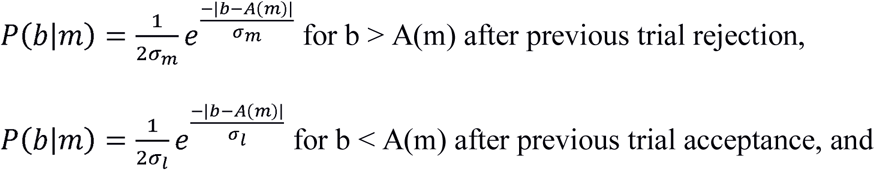

for the rest of (rare) cases, where *P(b*|*m)* is the Laplace distribution of bids *b* for market type *m*, and *σ_m_*, *σ_l_*, and *σ_0_* are parameters proportional to the standard deviation of the Laplace distribution. This captures the intuition that the tail above the preferred bid after rejections is fatter than the tail below the preferred bid after acceptances.

#### DL delta rule with asymmetric leptokurtic noise

This algorithm incorporates both the asymmetric leptokurtic policy distribution and the delta rule-based updating of the preferred bid. This was the best-fitting algorithm (Figure 2A, Table 1). It included an additional parameter *k* which accounted for a different proportion of trials with explorative (risky) versus exploitative (safe) bids.

**Figure 2.**
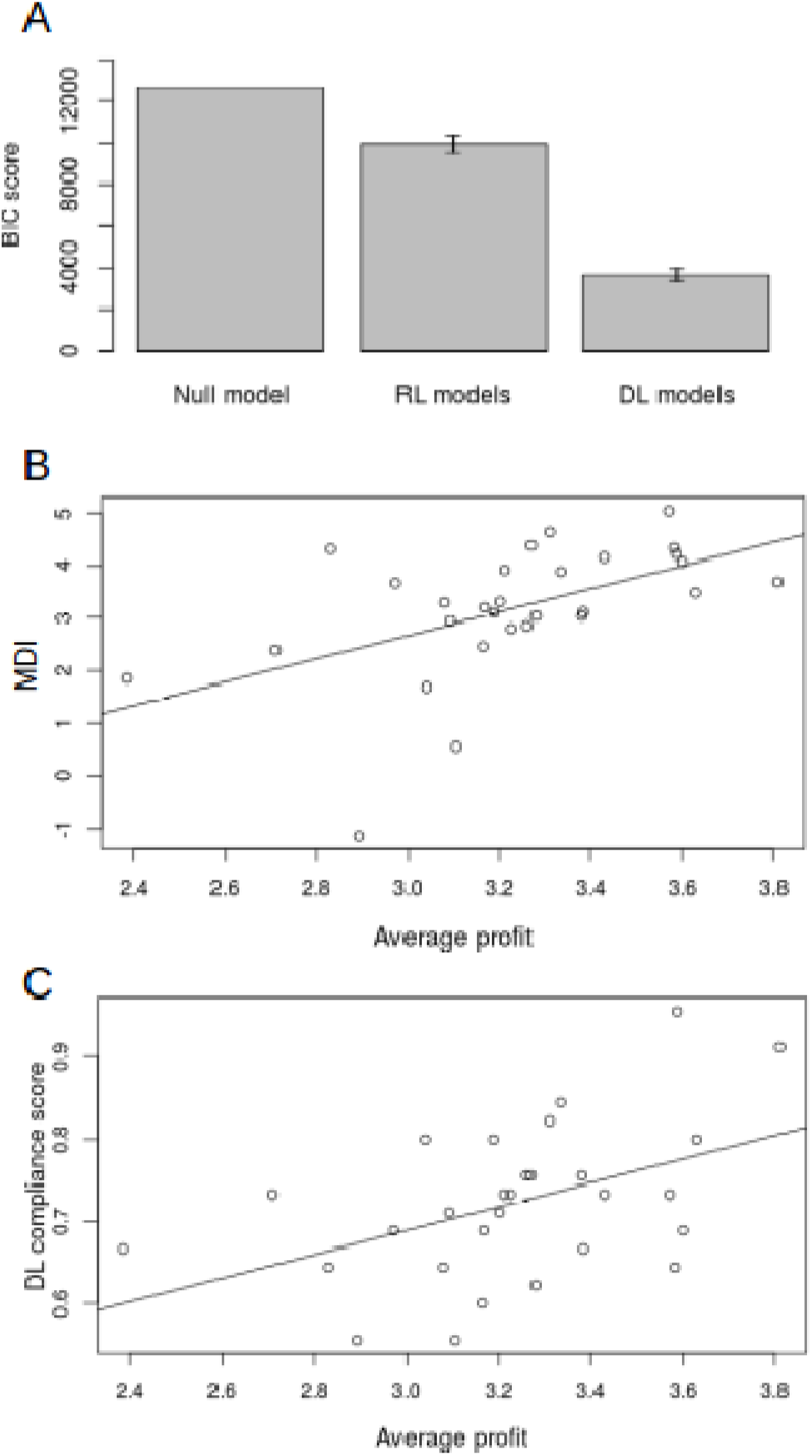
Algorithm fit scores and correlations with individual profits during the task. (A) BIC scores averaged within algorithm classes (DL: models 1-3, RL: models 4-6 in Table 1). (B) Correlation of market differentiation index with profits averaged across the whole task. The line slope correspons to a (Pearson’s product-moment) correlation coefficient of 0.524 (p=0.003). (C) Scatter plot of subjects’ DL compliance scores and profits averaged across the whole task. The line slope corresponds to a correlation coefficient of 0.466 (p=0.01). N=27. Error bars indicate 95% confidence intervals.

### Learning algorithms optimization and software

Following the usual approach in estimation problems with a small number of trials, a global objective function (the log-likelihood of aggregated data) was optimized with yoked parameters (fixed effects) across all subjects for all learning algorithms (Daw et al., 2006). This reduces parameter estimator variances at the cost of losing the ability to make between-subject parameters comparisons by pooling together between-subject and within-subject variability, but this is deemed to have little impact in the quality of the algorithm simulation predictions (Grinband et al., 2008). Given the scarcity of within-subject samples and the jagged geometry of the resulting objective functions, and that the random and fixed-effects analyses yielded largely consistent results (Table 1), we preferred this fixed effects comparison over the alternative of running the numerical optimizer for each subject individually in an objective function with multiple local extrema, which can lead to overfitting and bad performance of the numerical optimizer (but see Wilcox, 2005). For each algorithm agent, negative log-likelihood functions were constructed by making the agent play all 27 of the subject sessions. The log-likelihood function was

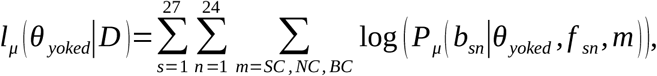

where *l_μ_* is the log-likelihood function for model *μ*, *θ_yoked_* is the parameters vector of model *μ* (for example, for naive RL *θ_yoked_*=(α,β)), and *P_μ_* is the likelihood of model *μ* choosing a specific bid *b* given parameters *θ* and feeback *f_sn_* in market type *m* for subject *s* and block number *n*. A numerical local-search optimizer was then run on each of the negative log-likelihood functions, and the found minima were used to recover the maximum likelihood parameter estimations. Bayesian Information Criterion (BIC) scores were derived from the negative log-likelihood values (Table 1).

To check for consistency, we also performed separate optimization routines for each subject objective function: *l*_*μ*,*s*_(*θ_s_*|*D_s_*), with individual free parameters *θ_s_* for subject *s*. The scarcity of data samples prevented convergence in some subjects, but converged instances yielded consistent BIC scores and parameter fits (Table 1).

Because subjects have 101 possible actions and they play only 60 times in all three market types, convergence of the model-free RL algorithms is troublesome when parameters are fitted individually, since values are updated sparsely and rarely, and often the game ends without sampling all possible states or actions. This is a problem for algorithm fitting, and in particular estimating 101 initial action-values depletes all useful degrees of freedom during optimization. Therefore, either we simplified the initial action-values using a three-parameter (as opposed to 101) Beta distribution or else we simply used the first round bids as initial conditions.

Data were processed with code written in Python with the scientific computing packages Numpy (RRID:SCR_008633), Scipy (RRID:SCR_008058), Matplotlib (RRID:SCR_008624), and Pandas. Purpose-specific code was written to define the maximum likelihood functions used to estimate the parameters of the learning algorithms. The numerical optimizer employed was a bound-constrained version of the Broyden-Fletcher-Goldfarb-Shannon algorithm, a local search technique which approximates local curvature. This algorithm is an implementation of a constrained optimizer of multivariate scalar functions belonging to the Python package Scipy. This optimizer was combined with a basin-hopping heuristic (scipy.optimize.basinhopping) with at least ten “hops” to offset the probability that the optimizer would converge into a local minimum due to the jagged geometry of the log-likelihood function.

### fMRI data collection and analysis

#### Data acquisition

The fMRI data were obtained using ascending interleaved slice acquisition with gradient echo T2^∗^–weighted echo-planar imaging (EPI) sequence in a 3T Magnetom Verio equipped with a 32-channel head coil (Siemens; Erlangen, Germany). Scanning protocol parameters were as follows: TE=30 ms; flip angle=80°; TR=2280 m; slice thickness=3 mm; no gap; slice matrix=64×64; number of axial slices=35; FoV=192 mm; Voxel resolution=3×3×3.7 mm.

High-resolution structural MRI data acquisition used a T_1_–weighted MP-RAGE sequence. Parameters were as follows: TE=2.47 ms; flip angle=9°; TR=1900 ms; slice thickness=0.5 mm; slice matrix=512×512×176; number of slices=176; FoV=256 mm; Voxel resolution=0.508×0.508×1 mm. These data were used for anatomical localization. A corrective routine aimed at counteracting susceptibility angled through the slice plane (z-shimming) was performed by the scanner. The slice angle was tilted a negative 30° with respect to the anterior commissure–posterior commissure axis in the sagittal plane to reduce the unaccounted spatial components of the susceptibility gradients (Weiskopf et al., 2006) and because this allows for better acquisition of the orbitofrontal cortex (Deichmann et al., 2003). The number of volumes acquired was on average 1,263, corresponding to a duration of approximately 48 min.

#### Sample size

Although median (reward-related) effects sizes in the striatum for gambling have been reported to have a Cohen’s d of only 0.4 (Poldrack et al., 2017), and our study was in theory only sufficiently powered to detect effects greater than 0.76 (based on sample size of 27 subjects and the analysis of Poldrack et al. (2017), using the distribution of local maxima of Cheng and Schwartzman (2015)), we found significant population level striatal signals with 27 subjects.

#### Preprocessing

Images were processed using SPM12 (Wellcome Department of Imaging Neuroscience, Institute of Neurology, London, UK). Preprocessing of T2^∗^–weighted volumes consisted of rigid-body model realignment within each session to a mean volume for head-motion correction, unwarping of the residual variance using the field map, slice-timing correction centered at TR/2, bias-field correction, coregistration of T2^∗^–weighted volumes to the corresponding structural image (T1–weighted volume) and segmentation and spatial normalization to a standard T2^∗^–weighted template (Montreal Neurological Institute, MNI) for group analysis, spatial smoothing with an 8 mm Gaussian kernel, and high-pass temporal (128s) filtering. Fieldmaps were acquired using a dual echo 2D gradient echo sequence with echoes at 5.19 and 7.65 ms, and repetition time of 444 ms, and then used with the SPM FieldMap toolbox to correct EPIs for unwanted dropout due to variations in spatial magnetic susceptibility (Weiskopf et al., 2006; Jezzard & Balaban, 1995).

#### GLM analysis

Eight event-related regressors (delta sticks) were used to model the onset of the MARKET stage (MARKETxSC, MARKETxNC, MARKETxBC), LOTTERY outcome stage (for won and lost lotteries), CHOICE stage and OUTCOME stage (ACCEPTED and REJECTED). In addition, five parametrically modulated delta sticks were constructed: three for all stages of the task using the *preferred bid value* (PBV=10-PB): MARKET_PBV, LOTTERY_PBV, CHOICE_PBV; one for the pseudo-RPE signal at outcome (OUTCOME_pseudo-RPE) *based on the best*-*fitting DL algorithm*; and one for the DS signal (OUTCOME_DS, consisting of +1 for positive RPEs and −1 for negative RPEs). Importantly, according to DL, the variables tracking currently estimated action values were not conventional expected values, but rather an estimation of the value of the maximum reward obtainable (PBV). Computing an expectation over a probability distribution of values associated with actions is not possible in a DL algorithm because there is no action value function over which a measure can be integrated. Thus, PBVs should be interpreted as a rough equivalent of the conventional expected values of RL algorithms. Both parametrically modulated and non-modulated stimuli onset markers were convolved (first order expansion) with the canonical hemodynamic response function (HRF) implemented in SPM12 and entered into a general linear model (GLM). The motion parameters output from the preprocessing realignment routine were added to the design matrix as covariates to account for residual head-motion effects.

In a separate analysis, two additional GLM regression matrices with three regressors were constructed with the stimulus onset marker OUTCOME and the mutually orthogonalized OUTCOME_DS and OUTCOME_RPE parametrically modulated regressors.

ROI activity in basal ganglia and PPC was extracted with the SPM extension MarsBar (Brett et al., 2002). Masks consisted of 8-mm spheres with center in-peak cluster of activity associated to preferred bid value in PPC (MNI coordinates [+-47,-48,52]), and manually delineated anatomical subdivisions of basal ganglia were used as in Palminteri et al. (2015), in both cases with their contralateral homologues. Specifically, (beta) coefficient estimates were calculated by averaging over the coefficients of all voxels within their ROIs separately for each subject. These individual estimates were passed to the group-level analysis, where the final coefficient values and their confidence intervals were calculated using the summary statistics approach.

#### fMRI statistics

Temporal serial correlations in fMRI data were removed with the (SPM12) autoregressive AR(1) model to satisfy the parameter estimation routine (restricted maximum likelihood) assumptions. Each subject design matrix was fitted individually, and the resulting regression coefficients were taken to a random effects group-level analysis. This analysis was performed by means of a summary statistics approach (Holmes & Friston, 1998). All reported fMRI statistics come from the group level.

Most decision-making studies model brain activity lasting less than 4 s with delta sticks, but studies have shown that this activity often lasts until the motor response (Grindband et al., 2008). Therefore, to ensure that such effects were not being ignored, we repeated the same analysis but with boxcar-shaped regressors functions instead of delta sticks. We found no significant additional effects.

Activations were also reported for regions of interest (ROI): striatum (Palminteri et al., 2015), orbitofrontal cortex, frontopolar and dorsolateral prefrontal cortex, anterior cingulate cortex, and medial prefrontal cortex, and temporo-parietal junction (Tzourio-Mazoyer et al., 2002). Activations were reported at a voxel-level threshold of p<0.05 after family-wise error rate (FWER) correction outside ROIs and for the learning signals (DS and pseudo-RPE) in the striatum because these activations were much stronger than any other. Brain regions are displayed on a standard MNI template. All clusters from all figures are listed in Tables 2, 3, and 4.

**Table 2.** Neural activity related to market type recognition and expected value (Figure 3).

| Contrast (Figure) | Region | Cluster p-value<br>FWE-corrected | Cluster<br>extent k | z-score | Peak voxel<br>p-value | MNI<br>(x, y, z) |
| --- | --- | --- | --- | --- | --- | --- |
| MARKETxBC vs MARKETxNC<br>(Figure 3A Left) | Left SPL | 0.085 | 43 | 4.33 | <0.001 | -33 -46 48 |
|  | Right SPL | 0.044 | 53 | 3.87 | <0.001 | 36 -46 60 |
|  | Right ANG |  |  | 3.44 | <0.001 | 39 -46 45 |
| MARKETxSC vs MARKETxNC<br>(Figure 3A Right) | Left SPL | 0.818 | 9 | 3.32 | <0.001 | -33 -52 48 |
| CHOICE_PBV<br>(Figure 3B) | Left SPL | 0.630 | 15 | 3.49 | < 0.001 | -47 -48 52 |
| REJECTED vs ACCEPTED,<br>MDI-modulated,<br>group level (Figure 3C) | Right SFG | 0.031 | 76 | 4.15 | <0.001 | 21 59 19 |
|  | Left SFG | 0.125 | 47 | 3.83 | <0.001 | -24 53 23 |
|  | Right MFC | 0.582 | 17 | 3.79 | <0.001 | 6 29 -14 |
|  | Right ANG | 0.301 | 30 | 3.66 | <0.001 | 60 -52 23 |
|  | Right TrIFG | 0.258 | 33 | 3.61 | <0.001 | 54 32 4 |
|  | Left MSFG | 0.528 | 19 | 3.56 | <0.001 | -3 50 4 |

Here and below all voxels are significant at least at $p < 0.001$ , uncorrected for the whole brain. Cluster-level inference alone may yield inflated false positives (Eklund et al., 2016; Flandin and Friston, 2017). x, y, z: stereotactic coordinates of the MNI template. Atlas labels were provided by Neuromorphometrics, Inc. MSFG: superior frontal gyrus medial segment; TrIFG: triangular part of the inferior frontal gyrus; ANG: angular gyrus; MFC: medial frontal cortex; SFG: superior frontal gyrus; SPL: superior parietal lobule; OCP: occipital pole; MFG: middle frontal gyrus; STG: superior temporal gyrus; MorG: medial orbital gyrus; CblExt: cerebellum exterior; AIns: anterior insula; NAcc: accumbens area

**Table 3.** Neural activity coding error signals pseudo-RPE and DS (Figure 4).

| Contrast (Figure) | Region | Cluster p-value<br>FWE-corrected | Cluster<br>extent k | z-score | Peak voxel<br>p-value | MNI<br>(x, y, z) |
| --- | --- | --- | --- | --- | --- | --- |
| DS<br>(Figure 4A Left) | Left Putamen | <0.001 | 47 | 5.59 | <0.001 | -30 -10 8 |
|  | Right CblExt | <0.001 | 147 | 5.51 | <0.001 | 33 -58 -40 |
|  | Left MorG | <0.001 | 20 | 5.50 | <0.001 | -24 35 -18 |
|  | Right Putamen | <0.001 | 35 | 5.48 | <0.001 | 30 -10 4 |
|  | Left CblExt | <0.001 | 83 | 5.36 | <0.001 | -15 -52 -18 |
|  | Left Caudate | <0.001 | 16 | 5.34 | <0.001 | -24 -19 23 |
|  | Right Caudate | <0.001 | 51 | 5.33 | <0.001 | 24 -10 26 |
|  | Right Putamen |  |  | 5.20 | <0.001 | 24 14 0 |
|  | Right CblExt | 0.001 | 9 | 5.25 | <0.001 | 6 -70 -33 |
|  | Right OCP | 0.001 | 12 | 5.21 | <0.001 | 18 -100 8 |
|  | Left Caudate | <0.001 | 13 | 4.94 | <0.001 | -21 11 19 |
|  | Right SPL | 0.003 | 6 | 4.91 | <0.001 | 45 -43 60 |
| pseudo-RPE<br>(Figure 4A Center Left) | Right CblExt | <0.001 | 119 | 5.83 | <0.001 | 18 -67 -22 |
|  | Left OCP | <0.001 | 25 | 5.32 | <0.001 | -12 -103 4 |
|  | Right NAcc | <0.001 | 48 | 5.28 | <0.001 | 12 17 -11 |
|  | Right Putamen |  |  | 5.28 | <0.001 | 21 14 -11 |
|  | Right Putamen | <0.001 | 14 | 5.14 | <0.001 | 30 -13 8 |
|  | Left SMG | 0.003 | 7 | 5.11 | <0.001 | -57 -34 45 |
|  | Left MFG | 0.001 | 10 | 5.04 | <0.001 | -36 35 30 |
|  | Left MFG | 0.002 | 9 | 4.87 | <0.001 | -39 38 15 |
|  | Right OCP | 0.004 | 6 | 4.77 | <0.001 | 15 -100 11 |
|  | Left CblExt | 0.003 | 7 | 4.76 | <0.001 | -12 -52 -22 |
| ort-pseudo-RPE<br>(Figure 4A Center Right) | Left MFG | <0.001 | 197 | 4.23 | <0.001 | -24 20 63 |
|  | Right SPL | 0.315 | 29 | 3.93 | <0.001 | 27 -61 34 |
|  | Right MFG | 0.196 | 38 | 3.92 | <0.001 | 42 14 56 |
|  | Left SPL | 0.023 | 82 | 3.89 | <0.001 | -21 -46 45 |
|  | Right SFG | 0.283 | 31 | 3.71 | <0.001 | 27 14 63 |
|  | Right MFG | 0.501 | 20 | 3.63 | <0.001 | 36 38 30 |
|  | Right MFG | 0.924 | 5 | 3.56 | <0.001 | 48 41 26 |
|  | Right ACgG | 0.728 | 12 | 3.52 | <0.001 | 12 38 11 |
|  | Left Nacc | 0.609 | 16 | 3.50 | <0.001 | -9 8 -7 |
|  | Left Caudate | 0.788 | 10 | 3.42 | <0.001 | -15 -4 23 |
|  | Right MFG | 0.924 | 5 | 3.42 | <0.001 | 39 47 8 |
|  | Left ACgG | 0.924 | 5 | 3.25 | 0.001 | -3 32 -11 |
| ort-DS<br>(Figure 4A Right) | Left Caudate | 0.070 | 56 | 4.36 | <0.001 | -27 -7 26 |
|  | Left Putamen |  |  | 3.79 | <0.001 | -27 -10 8 |
|  | Right Caudate | 0.227 | 34 | 4.18 | <0.001 | 24 -10 26 |
|  | Right Putamen |  |  | 3.60 | <0.001 | 27 -10 11 |
|  | Right STG | 0.057 | 60 | 4.07 | <0.001 | 57 -28 8 |
|  | Right Caudate | 0.543 | 18 | 3.97 | <0.001 | 21 20 15 |

**Table 4.** Neural activity during OUTCOME stage associated with follow-up bid increases (Figure 5).

| Contrast (Figure) | Region | Cluster p-value<br>FWE-corrected | Cluster<br>extent k | z-score | Peak voxel<br>p-value | MNI<br>(x, y, z) |
| --- | --- | --- | --- | --- | --- | --- |
| ACCEPTED<br>bid increase-modulated<br>(Figure 5A) | Right Caudate | 0.515 | 16 | 4.22 | <0.001 | 18 5 19 |
|  | Right Putamen | 0.020 | 59 | 4.18 | <0.001 | 18 8 -11 |
|  | Right AIns |  |  | 3.57 | <0.001 | 33 11 -18 |
|  | Left MFG | 0.764 | 10 | 3.92 | <0.001 | -33 56 19 |
|  | Left MFG | 0.035 | 51 | 3.87 | <0.001 | -30 41 34 |
|  | Right SMG | 0.047 | 47 | 3.85 | <0.001 | 63 -34 19 |
|  | Left Putamen | 0.202 | 28 | 3.79 | <0.001 | -21 8 -7 |
|  | Right SFG | 0.917 | 6 | 3.40 | <0.001 | 24 44 26 |
|  | Left MSFG | 0.806 | 9 | 3.34 | <0.001 | -9 50 0 |
| REJECTED<br>bid increase-modulated<br>(Figure 5B) | Right Putamen | 0.818 | 9 | 3.59 | <0.001 | 24 14 -3 |

We also investigated correlations between neural data and model proxy variables to localize potential brain regions involved in the computation of the economic transactions on a trial-by-trial basis. We derived a time series of expected value and prediction error signals from each of the bidding algorithms by simulating bidding agents with each algorithm and pitting them against the same sequences of stimuli that the human subjects played against. The dataset comprising all the sequences for all subjects was used. Then, we fitted fMRI data to the learning algorithm variables informed by plausible assumptions on the strategies used by players to maximize profit, and we selected the best algorithms based on BIC scores.

We standardized all algorithm proxy variables (PBV, pseudo-RPE, DS) as z-scores across subjects before entering them as parametric regressors in the design matrix. In the group-level analysis, we used this analysis to link between-subject differences to significant activations (Haruno et al., 2004).

Finally, a neural model comparison routine based on a SPM Bayesian model selection module was performed on anatomical ROIs encompassing striatum and inferior posterior parietal cortex. To assess the goodness of fit of both DL and RL algorithms to neural activity, we defined GLMs in OUTCOME, including either DS or RPE parametric modulators, respectively, and then estimated them using Bayesian statistics, which provided a measure of the evidence of the model for each subject. Log evidence was then fed to a BMS random-effect analysis (Palminteri et al., 2015; Stephan et al., 2009), which computed the *exceedance probability* of each GLM within the anatomical mask.

## Results

### Behavior across market types indicates heuristic (DL) learning of valuation

Overall, subjects successfully performed the double auction task under all types of social competition (72.47% of successful transactions). Transaction rates per market type were 92.44% (869/940) in SC, 74.68% (702/940) in NC, and 50.26% (472/939) in BC market.

To estimate subjects’ beliefs about their human opponents and each market type prior to learning, we compared the bids in the first trial of each experimental session. On average, subjects bid 4.96, 5.13, and 6.55 monetary units (MUs) in the SC, NC, and BC markets, respectively. A one-way ANOVA test rejected the hypothesis that first mean bids were equal: F(2,137)=18.93, p=6^∗^10^−8^. Thus, subjects discriminated among market types already before the beginning of the task. Reaction times (RT) did not differ significantly across market types: 11.2±3.6s, 11.1±3.8s, and 11.8±3.8s for SC, NC, and BC, respectively.

Next, we wanted to know how the bids and bid-adjustments evolved over time and across markets. We tracked the evolution of subjects’ bid choices in each market (Figure 1B) by fitting a linear mixed-effects model with random intercepts. Subjects gradually decreased bids in SC (beta=−0.027, t(588)=−4.44, p=5.4^∗^10^−6^) and increased bids in BC (beta=0.086, t(587)=14.264, p=4^∗^10^−40^), whereas in NC, we found a weakly significant evolution of bids (beta=−0.009, t(588)=−2.01, p=0.045). Notice that the decreases in SC and increases in BC are not symmetric: subjects tended to increase the bids much more than decreasing them.

We reasoned that bid changes should depend directly on the subjects learning their success or failure in the previous bid they made. Hence, to inquire into the potential causes of bid evolution, we examined the effect of the previous trial outcome on the current bid. We tracked, on a trial-by-trial basis, the bid increments from one trial to the next within a given market type (Figure 1C). The distribution of these bid increments conditioned on the outcome of the previous trial displayed a skewed shape, with opposite skewness for the previous-trial -accept and -reject bids. Such distribution can be roughly sketched as an asymmetric accept-down/reject-up rule or win-stay/lose-shift strategies (Nowak & Sigmund, 1993). Furthermore, the distributions of bid increments were qualitatively invariant across all market conditions, suggesting that the trial-by-trial learning rule underlying bid adjustments is independent of the market type. Therefore, we reasoned that the subjects’ market-dependent bidding trends must be attributed largely to the opponents’ behavior. This supports a view where the subjects’ bid learning strategy (or algorithm) does not change from market-type to market type; yet the subjects explicitly recognize which market condition they are. This is indeed suggested by data in Figure 1B showing that the bids are rapidly rescaled between the different market types. We thus inquired what formal learning algorithm could best account for the learning behavior and the evolution of bids (irrespective of the market type): conventional model-free RL algorithms or model-based algorithms that take into account the structure of the task (see below).

Finally, we examined whether subjects’ ability to bid successfully was related to how well they learned to identify the different market conditions. To get a coarse index of the degree to which subjects distinguished between the three market types, we devised the market discrimination index (MDI), calculated as the difference between the mean bid chosen over all trials for BC and SC conditions. Buyers who better distinguished market types, as assessed by the MDI, were more likely to receive higher profits (Figure 2B). Indeed, we found a significant correlation between profit earned and the MDI (r=0.52, Pearson’s product-moment correlation, t=3.20, df=27, p=0.003497, 95%CI=[0.1955, 0.7473]). Thus, in our task better market discrimination is associated on average with higher profit.

The above results gave us a hint that the observed behavior may be accounted for by a DL algorithm of bid learning, where bids are nudged up or down depending on previous outcome. Importantly, DL requires a model of the “action (bid) space” to account for the directionality of bid adjustments. We also note that the traditional reinforcement learning schemes and DL differ in the learning signals they use to update decision-making variables: a continuous reward prediction error (RPE) for RL and a binary error signal we denote by directional signature (DS) for DL (see Methods for details). In order to test our hunch that DL is used to learn bids in our task, we proceeded to test which DL or (and) RL algorithms could best explain the observed behavior.

### Adaptive learning algorithm fits and model selection

We fitted six adaptive learning algorithms to the behavioral data. All DL algorithms fitted better than RL algorithms (Figure 2A). As we expected, the RL algorithms failed to explain the bid evolution in all market types. We believe this was in part due to a difference in the efficiency of the two algorithm classes. The RL algorithms require a large dataset to learn action values to the point where they start being operationally useful. Since our subjects learned to bid successfully in the limited number of played trials, we argue that DL is the more efficient and appropriate learning strategy for the task we consider.

Across all subjects, 74.99% (1586/2115) of the trials matched the behavioral predictions of the best DL algorithm. Conditioned on the outcome of the previous trial of the same market type, subjects behaved according to the DL algorithm in 76.26% (453/594) and 79.73% (1133/1421) of trials when their bids were rejected and accepted, respectively. Importantly, subjects with a higher *DL*-*compliance score* (the fraction of trials where they behaved according to DL) were more likely to receive higher profits (Figure 2C). We found a significant between-subjects correlation between the profit earned and the proportion of trials compliant with DL (r=0.47, Pearson’s product-moment correlation, t=2.74, df=27, p=0.01078, 95%CI=[0.1204, 0.7113]). To confirm this we took the best-fitting DL and the best-fitting RL models and simulated their bidding against the same prerecorded opponents as the subjects. Only the DL agent’s bid evolution resembled the human one, with progressive increase in the SC bids and relative invariance of the NC and SC bids (not shown). Next, we proceeded to determine the neural underpinnings of repeated bidding learning.

### Fronto-parietal cortical activity associated with recognition of the different market types

To identify the brain regions associated with subjects recognizing the different market types, we analyzed the neural activity during the MARKET stage of the task, which informs subjects about the market type at the beginning of each trial. We found that neural activity in the posterior parietal cortex (PPC) increased when subjects entered the competitive BC and SC markets (Figure 3A, Table 2) as compared to NC. The effect remained significant when the expected reward based on the preferred bid was regressed out, ruling out that it was a value-related activation. The other pairwise subtraction contrasts between market types revealed no significant differences in activity.

**Figure 3.**
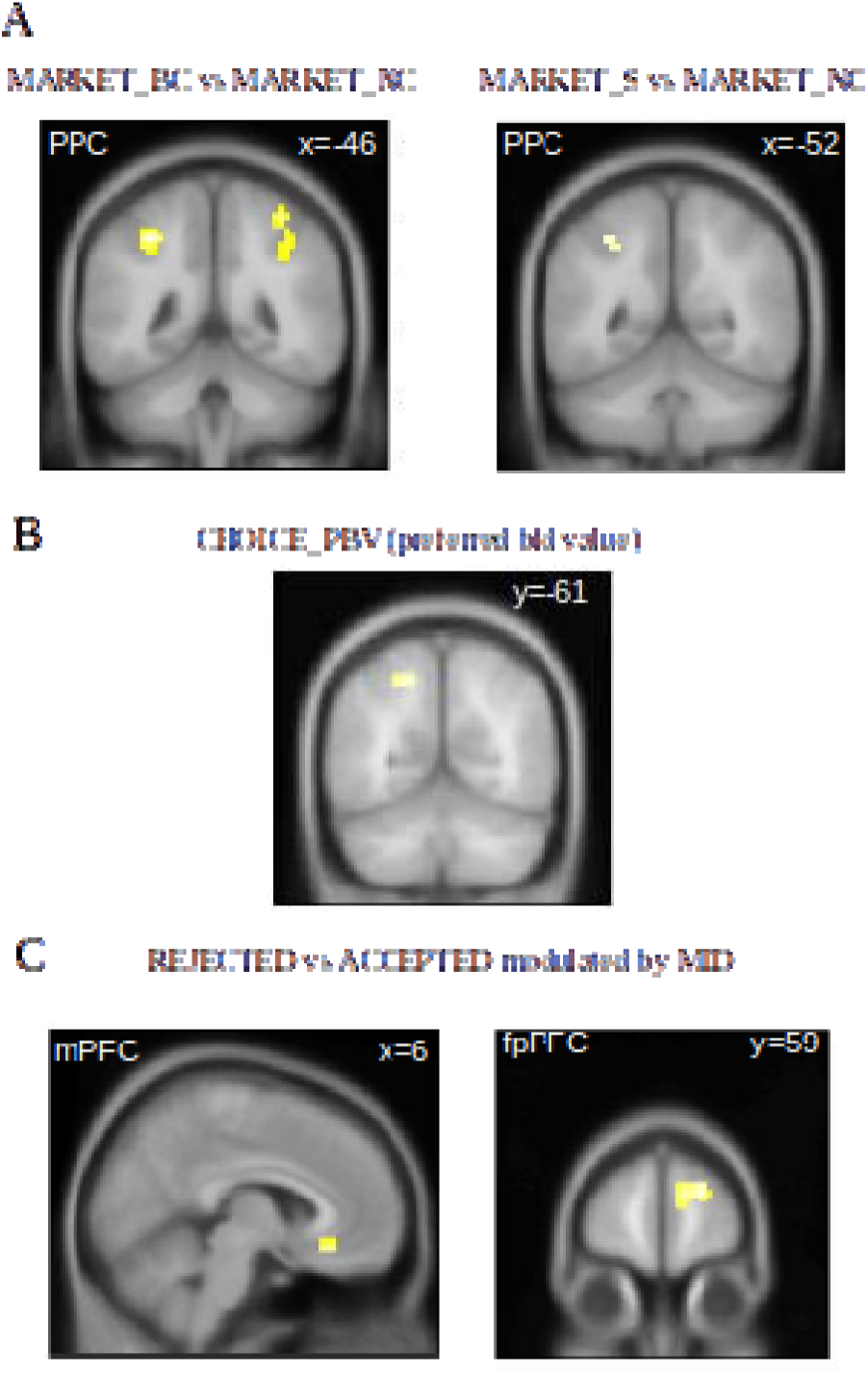
Neural activity related to market type recognition and expected value. (A) Left: stronger superior parietal cortex activity in BC as compared to NC condition during market entrance (MARKET_BC vs MARKET_NC). Right: stronger left superior parietal cortex activity in SC market as compared to NC market during market entrance (MARKET_SC vs MARKET_NC). (B) Activation reflecting modulation by the preferred bid during bid choice (CHOICE_PBV). (C) Feedback processing-related activity (outcome stage, REJECTED vs ACCEPTED) modulated by individual differences in market differentiation index in the right medial frontal cortex (C Left) and frontopolar cortex (C Right). Clusters are listed in Table 2

To further investigate neural activity underlying the recognition of the different market types, we used the MDI as a covariate in the group-level analysis. The between-subject differences were manifested only in the prefrontal activity during processing of outcomes (OUTCOME stage, Figure 3B), specifically in a region bridging the bilateral medial frontal and superior frontal gyrus, adjacent to the frontopolar prefrontal cortex (fpPFC), and in mPFC (Figure 3C). Thus, fronto-parietal activity was associated with the recognition of market types.

### Posterior-parietal cortex activity associated with the internal representation of bid space

To find brain areas whose activity encoded an internal representation of bid space, we used the preferred bids provided by the fitted DL algorithm as a covariate regressor at the CHOICE stage. We found a significant activity modulation in the PPC (Figure 3B). This indicates that learned preferred bids are encoded in the PPC. Bids are real numbers, and their representation in the PPC is compatible with previous studies showing evidence for encoding of a number line in PPC (Dehaene et al., 2003). Moreover, the PPC region associated to the preferred bid value was also strongly modulated by both pseudo-RPE and DS signals (Figure 4).

**Figure 4.**
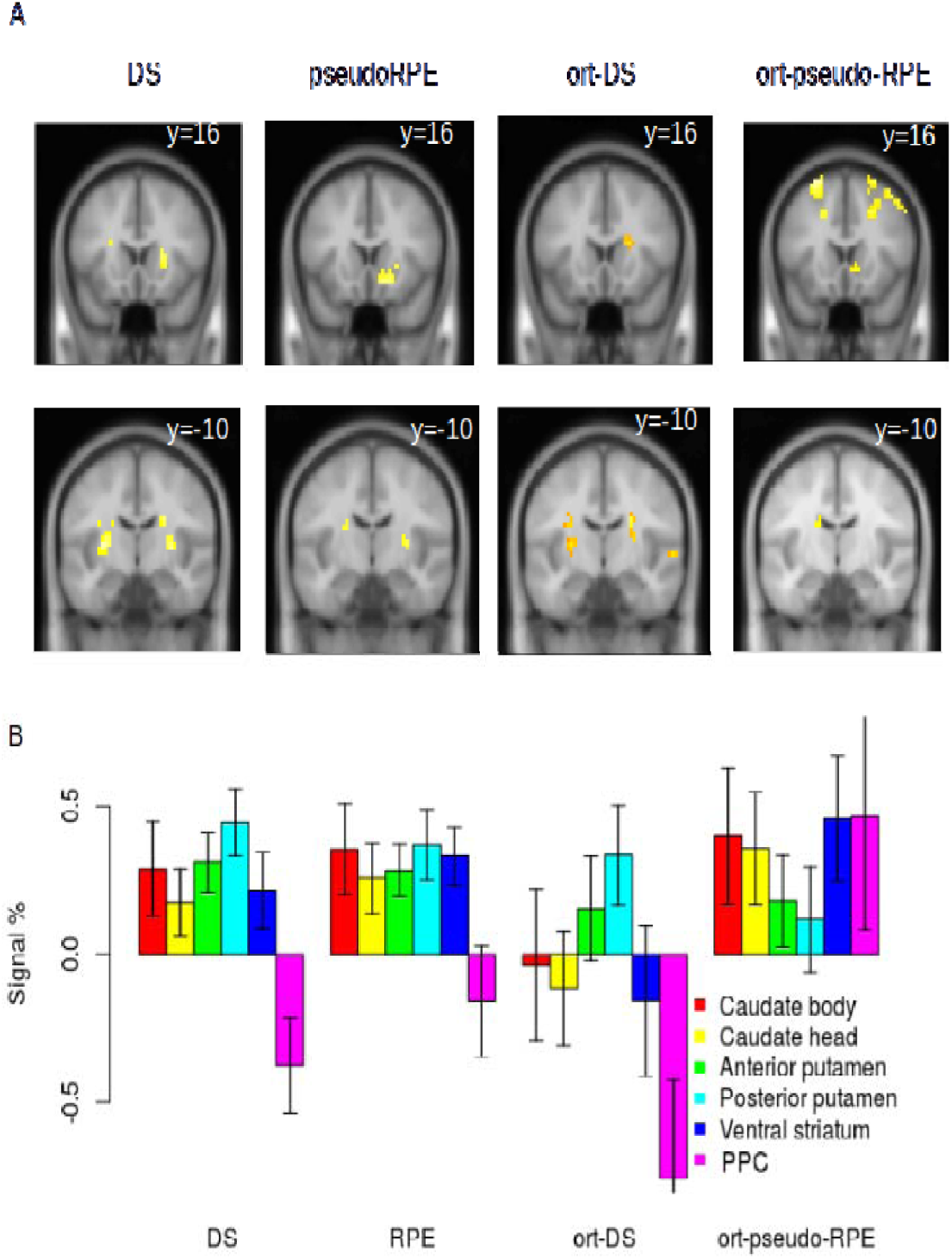
Neural correlates of pseudo-RPE and DS signals *based on the best*-*fitting DL algorithm* in anterior putamen and nucleus accumbens area and posterior putamen during OUTCOME. (A) Correlated activity in the anterior (y=16) and posterior (y=−10) putamen was stronger for pseudo-RPE and DS, respectively, during feedback. From left to right columns: pseudo-RPE (p<0.05, FWER), DS (p<0.05, FWER), pseudo-RPE orthogonalized with respect to DS (p<0.001, unc), and DS orthogonalized with respect to pseudo-RPE (p<0.001, unc). (B) Barchart of signal estimation (in grand mean percentage) by brain region. Signals were averaged within anatomical ROIs for basal ga glia (Palminteri et al., 2015) and on an 8mm sphere in PPC. oDS and oRPE correspond to DS and pseudo-RPE signals after being orthogonalized with respect to each other, respectively. Clusters are listed in Table 3.

### Striatal activity associated with trial-by-trial adaptive learning

In order to identify the neuronal representation of the learning algorithms used, we compared the explanatory power of RL and DL algorithms over the neural activity in the two areas most relevant to the task: striatum and PPC. We calculated the *exceedance probability* (Stephan et al., 2009) for each algorithm, given the brain imaging data gathered from all subjects. The exceedance probability was calculated using Bayesian model comparison of GLMs regressing the learning signals, DS for DL and pseudo-RPE (the RPE based on the accepted preferred bids of the DL algorithm, see below) for RL. The analysis confirmed the explanatory power of the DL algorithm to be stronger than that of the RL algorithms: the P_exc_(DL)=0.9533 > P_exc_(RL)=0.0467. This yields a Bayes factor above 19, which indicates clearly strong evidence (Kass & Raftery, 1995) in favor of DL.

Therefore, we used the variables provided by the best-fitting DL algorithm to search for neural correlates of the outcome evaluation and learning during the CHOICE and OUTCOME stages. In particular, we asked whether DL and RL neural learning signals could be distinguished. We reasoned that it is unsound to search for correlates of the variables extracted from the ill-fitting RL algorithms. Therefore, we instead compared RPE and DS signals by using the best-fitting DL algorithm and calculating RPEs based on the reward expected from accepted preferred bids, which we call pseudo-RPE. We then performed a whole-brain analysis for the OUTCOME stage and compared DS and pseudo-RPE.

Neural correlates of both DS and pseudo-RPE were found in the striatum (Figure 4). Because DS and pseudo-RPE are highly correlated, we orthogonalized both regressors with respect to each other: *ort*-*pseudo*-*RPE (pseudo*-*RPE orthogonalized with respect to DS)*, and *ort*-*DS (DS orthogonalized with respect to pseudo*-*RPE)*. Interestingly, *ort*-*DS* related activity was found primarily in the posterior putamen whereas *ort*-*pseudo*-*RPE* strongly modulated activity of the caudate and ventral striatum (Figure 4). This is in line with previous studies reporting that neurons in the caudate nucleus could play a role in transforming expected reward into a spatially selective behavior (Gold, 2003; Kawagoe et al., 1998; Lauwereyns et al., 2002).

Our results indicate that both DS and RPE signals are encoded in the striatum but in anatomically dissociated areas; anterior and ventral regions encode an RPE learning signal, whereas the dorsal and posterior regions encode a binary DS learning signal. We further explored averaged signals within anatomical ROIs. A two-way ANOVA (regions: [posterior striatum, anterior striatum], learning signal: [ort-DS, ort-pseudo-RPE]) yielded a significant interaction (p=0.0012; F=11.08, df=1). Although both signals are represented concomitantly, computational algorithm fits suggest that DS is the predominant learning signal.

Finally, we examined the relationship between learning-related neural activity during OUTCOME and the behavioral adjustments. We computed a parametrical regressor modulated by the size of the subsequent adjustments of bids (the bid in the next trial of the same market type minus the bid in the current trial). Given that subjects after the *accepted* trials usually repeated or sometimes decreased their bids, the activity of the dorsolateral prefrontal cortex (dlPFC) and the ventral striatum in *accepted* trials was associated with subsequent bid repetition (Figure 5A). After the *rejected* trials, subjects most often increased or (less frequently) repeated the bid; activity of the right putamen during *rejected* trials was associated with subsequent bid increase (Figure 5B). Thus, neural activity in the dlPFC and striatum correlated with bid adjustments.

**Figure 5.**
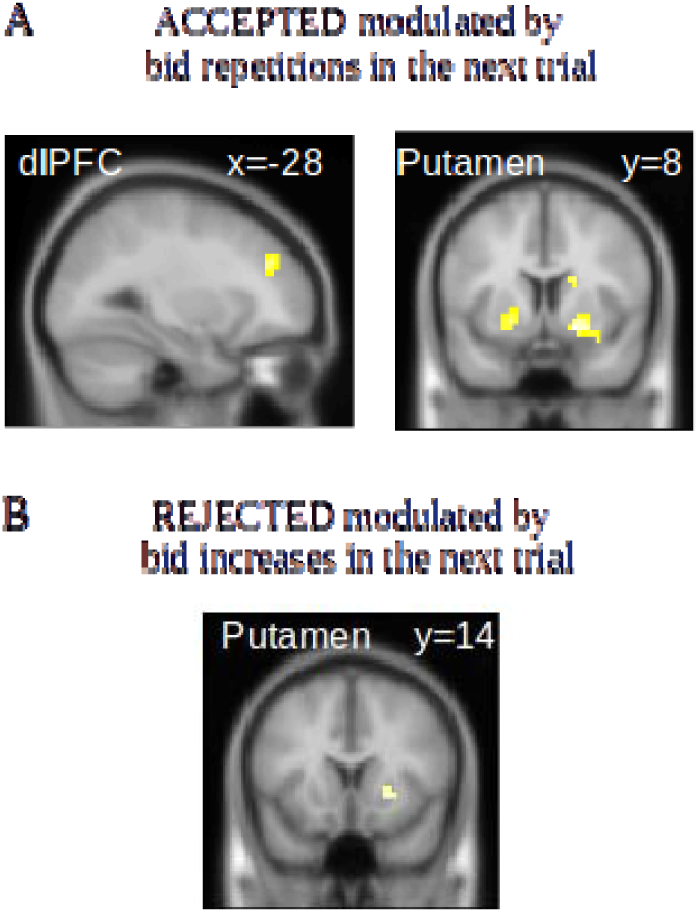
(A) Neural activity during positive feedback (ACCEPTED) in dlPFC (Left) and striatal (Right) areas that was significantly modulated by bid increases in the next trial of the same market type. (B) Neural activity during negative feedback (REJECTED) in putamen that was significantly modulated by bid increases in the next trial of the same market type. Clusters are listed in Table 4

## Discussion

We investigated the neural underpinnings of learning to bid in double auctions. We found that buyers learned to choose bids using an effective decision-making heuristic consisting of directional adjustments contingent on the previous trial outcome. As opposed to model-free reinforcement learning, directional learning postulates the existence of *a priori* knowledge about the structure of the task. Namely, DL assumes that the action values of bids bear an order relationship; it and stores and updates the value of the preferred bid on an internal number line. Therefore, DL naturally fits market and auction decisions in which prices or quantities are the main strategic variables. Although one could object that DL and RL are intimately related, a crucial aspect distinguishes them: unlike RL, DL does not learn an explicit value function spanning all actions, but only a single preferred action.

Analysis of the first bids in each market type revealed that subjects discriminated among the market types already at the beginning of the game. Although subjects underestimated the effect of social competition in the different market types, they gradually learned to optimize their bidding decisions. Indeed, the learning curve for each market type exhibited an incomplete convergence toward the strict Nash equilibrium predicted for perfectly rational agents. Importantly, the fact that the RTs did not differ across the market types suggests that the differences of learning curves in three markets were not confounded by cognitive effort differences.

Since numerous fMRI studies have demonstrated neural correlates of RPE in the striatum (e.g. Haruno et al., 2006; O’Doherty et al., 2003; van den Bos et al., 2013), we examined in detail pseudo-RPE and DS-related activity within this region. We found that the pseudo-RPE signal was observed in the anterior and ventral striatal areas, whereas the DS signal was represented in the dorsal posterior striatal areas, particularly in the posterior putamen. According to the Bayesian model comparison analysis, the variability of the striatal activity was explained by DL better than by RL, supporting the pertinence of DL-based bidding. This finding concurs with previous suggestions that neural learning signals in motivated decision-making are not necessarily always RPE-like (Behrens et al., 2008, supplement), and that a region of striatum is involved in learning stimulus–response associations and action selection (Jessup & O’Doherty, 2011). Although the coexistence of these complementary yet exclusive value signals is not exceptional (Lebreton et al., 2013), the reason underlying the coexistence of DS and pseudo-RPE signals in the striatum is unclear, since only DS explains the behavior of participants. Although these learning signals are difficult to decorrelate, a follow-up study could clarify their relationship, in particular, whether these signals could be partially ancillary to bidding behavior and be part of a hybrid DL-RPE architecture.

Intriguingly, in a correlation analysis of feedback processing related neural activity with the next trial bid adjustment, we also found that both dlPFC and striatum activity were associated with bid increase or repetition in the next trial *regardless* of whether the bid was previously accepted or rejected (Figure 5A). We may posit that activity of the dlPFC subserves a cognitive control mechanism for tracking the preferred bid, and concomitantly striatal activity has a role in increasing the value of the currently preferred bid. This parallels the previously reported role of the dorsal striatum in updating action values (Balleine et al., 2007; Haruno et al., 2004; Lauwereyns et al., 2002; Palminteri et al., 2012) and the parametric working memory encoding in the PFC reported by Romo et al. (1999). Significant activity predicting bid adjustments after rejection was also present in the putamen when subjects’ bids were rejected. To account for the role of the striatum in updating bids instead of values, we speculate that because the task revolves consistently around the bid choice, the reference magnitude for updating values was not the expected reward, but the preferred bid, as suggested by the best-fitting DL algorithm. Although, to our knowledge, such function has not been attributed to the striatum in previous studies, it is plausible that at least some neuronal submodules could compute bids instead of expected rewards because in our task, the bid is the natural operational variable (bid size is the only quantity that needs to be tracked) and is perfectly anti-correlated with reward when accepted. The activity consistently associated with “nudging up” bids and a similar signal reported in the superior PPC (Figure 3B) lends support to this hypothesis.

The DL-based learning strategy requires a representation of an internal number line where the preferred bids are stored and actively updated. Our results indicate that this representation is implemented in the PPC (Figure 3A). Accordingly, Gläscher et al. (2010) also found neural signatures of model-based prediction errors analogous to DS in the PPC in a Markov decision task, and the superior PPC has been implicated in directing spatial attention to a representation of an internal number line (Hubbard et al., 2005). Moreover, we found activity associated with the preferred bid size in the left superior PPC, which has been also found to represent the relative value or probability of different actions (Sugrue et al., 2005). Thus, during bidding, activity of the superior PPC could not only modulate attention to the internal number line, but also contribute to decision-making. Other neuroimaging studies show that the activities of the superior PPC contribute to working memory (Koenigs et al., 2009), arithmetic facts (Dehaene et al., 2004; Pesenti et al., 2000), and quick value-based decision-making (Jocham et al., 2014). Altogether, the superior PPC could participate in a calculation and representation of the preferred bid that is transmitted to motor areas to execute appropriate motor commands.

The ability to recognize market types is also critical for successful bidding. At the beginning of each trial, activity in the bilateral superior PPC was stronger in trials with higher social competition (SC and BC; Figure 3A). This activation could reflect neural activity monitoring the competitiveness in the current trial or retrieving relevant information (Vilberg & Rugg, 2008) about the current market type (i.e. the preferred bid). Activity in the superior PPC has been previously implicated in the processing of numerical information needed for the forthcoming motor selection (Sawamura et al., 2002). Thus, the PPC could set bargaining decisions into the appropriate social competition context by associating the specific market type with its associated DL-learned preferred bid. Therefore, successful bidding could be subserved by the same computational processes underlying simple arithmetical calculations (Dehaene et al., 2004) and distance estimation. Between-subject differences associated with the ability to distinguish the different market types in our study affected the activity of the fpPFC and vmPFC. This might indicate that subjects who distinguished better among market types, besides earning more profits, exhibited stronger activation of the higher-order cognitive prefrontal areas associated with the appraisal of suitable models of the environment (Boorman et al., 2009) and mentalizing (Hampton et al., 2008; Coricelli & Nagel, 2009). Congruently with previous fMRI studies, fpPFC activity might be involved in appraising the behavior of opponents (Koechlin & Hyafil, 2007), whereas vmPFC activity might be involved in appraising the subject’s own valuation during feedback.

In this study, we used prerecorded opponent data, which could affect behavior through social preferences (van den Bos et al., 2008) and arguably may not allow us to disentangle precise market-based prior strategies from feedback-based learning. Although studies using live opponents (e.g. Carter et al., 2012) eschew this limitation, they cannot control well for variability induced by repeated mutual feedback, which was necessary in our study to control the bid variability in each market type. Further studies are needed to verify the role of feedback-based learning in double auctions.

In conclusion, while the buyers were bidding under different levels of supply and demand, their behavior was explained best by a simple learning heuristic. Between-subjects higher compliance with DL predicted higher payoffs. Our results suggest that the PPC encodes an internal representation of a bid space that serves as a model on top of which subjects adjust and select bids, and posterior striatal activity was associated with a simplified learning signal characterized by a binary learning signal. Individual differences during feedback associated with activity in the dlPFC and superior PPC indicate the critical role of at least a rudimentary prior knowledge of the structure of the task and the differences among market types. In summary, we suggest that a learning heuristic based on a binary learning signal distinct from the conventional RPE signal solves the problem of repeated bidding in double auctions. Showing the learning mechanisms underlying bidding under social competition, this study paves new pathways for the discovery of neural mechanisms engaged in competitive, dynamic, complex decisions.

## Acknowledgements

We thank Stefano Palminteri and anonymous reviewers for constructive comments on the previous versions of the manuscript. We thank Laurent Muller for his contribution to the study design. This study has been funded by the Russian Academic Excellence Project ‘5-100.’

## Notes

**Conflicts of interest** The authors declare no competing financial interests.

